# Allonursing (Co-BreeD): Its prevalence, functions and hypotheses across mammals and human cultures

**DOI:** 10.64898/2026.01.10.698751

**Authors:** Yitzchak Ben Mocha, Maike Woith, Sophie Scemama de Gialluly, Lucia Bruscagnin, Laura Pipper, Natalie Kestel, Sonny Agustin Bechayda, Nikhil Chaudhary, Christine M. K. Clarke, Paul A. Garber, Lee T. Gettler, Andrea Pilastro, Rubén Quintana, Andrew N. Radford, Heinz Richner, Stacy Rosenbaum, Eduardo S. A. Santos, Michael Griesser

## Abstract

Allonursing, where dependent young are nursed by females other than their mothers, has been observed across diverse mammalian species and human cultures. Yet, the proximate functions underlying this seemingly altruistic behaviour, and its benefits for the offspring, the mother, and the nursing allomother, remain poorly understood. To facilitate testing hypotheses concerning these questions, we compiled the to date largest allonursing dataset from wild populations (N = 101 species, including 133 populations and 5 human cultures). Using this dataset, we (i) map the taxonomic distribution of allonursing, thereby expanding the list of mammal species qualified as cooperative breeders by 17 additional species, (ii) differentiate cases of non-voluntary from voluntary allonursing and consequently reject the hypothesis that milk stealing explains regular allonursing in most species, (iii) quantify the within-population extent of allonursing to assess its significance across species and evaluate the energy/time saved for mothers, (iv) use phylogenetically controlled analyses to revisit the association between allonursing and polytocy (i.e., litter size >1), and provide an alternative explanation for this association. We conclude by proposing that allonursing has multiple functions, with the most empirically supported ones being provisioning and insurance against maternal loss. Finally, the presented *AlloNursing* dataset complements the Cooperative-Breeding Database (Co-BreeD) to further expand this integrative resource for cooperative breeding research.

## I. Introduction

Cooperative breeding is an offspring care system with the core characteristic of individuals providing care for offspring of their group members (i.e., alloparental care, see Glossary, (Brown, 1974; Clutton-Brock, 2006; Riehl, 2013). Since caregiving is costly (e.g., in terms of time and energy), investigating the proximate functions of alloparental care and its evolutionary drivers is essential for understanding the emergence and maintenance of cooperative breeding (Capilla-Lasheras, Wilson & Young, 2023; Olivier & Higginson, 2023).

Reliable identification of alloparental care is a prerequisite for classifying species as cooperative breeders, which in turn is required for comparative research on cooperative breeding (Cockburn, 2006; Schradin, 2017). Across bird and mammal species, offspring require different types of care, including nest/den construction and sanitation (Jing et al., 2009; Haddad et al., 2024), babysitting (Stanford, 1992; Whitehead, 1996), predator defence (Amat, Fraga & Arroyo, 1999; Santema & Clutton-Brock, 2013), transfer between locations (Stanford, 1992; Ben Mocha, Mundry & Pika, 2019), feeding (Ostreiher, 1997; Harding et al., 2004), or teaching (Thornton & McAuliffe, 2006; Griesser & Suzuki, 2017). Some of these behaviours, though, can be self-directed (e.g., chasing away a predator as self-defence) or directed towards own offspring (e.g., babysitting own offspring while others’ offspring are around). Under these circumstances, group members perform these de facto “caring” behaviours regardless of the presence of others’ offspring, and young benefit as a by-product of sociality rather than through cooperative offspring care *per se* (Jennions & Macdonald, 1994; Ben Mocha et al., 2023b). Distinguishing alloparental care from self-beneficial behaviours or parental care is therefore critical (to disentangle the evolution of sociality from that of alloparental care), but often methodologically challenging.

Allonursing is a form of alloparental care in which dependent mammalian young suckle from the teats of a female other than their mother (Glossary). Allonursing research offers valuable insights into cooperative breeding from both the methodological and biological perspectives. From a methodological perspective, allonursing can be used to confirm alloparental care reliably because it cannot be self-directed (i.e., a female cannot allonurse herself), and cases of non-voluntary nursing (e.g., when females are being suckled while asleep) can be identified (see Results and Discussion). From a biological perspective, allonursing exemplifies the alleged contradiction in cooperative breeding: why do females in some mammal species invest in others’ offspring? Nonetheless, the proximate functions underlying this behaviour, as well as its costs and benefits for the offspring, the mother, and the nursing allomother, are poorly understood (Figure 1).

**Figure 1.**
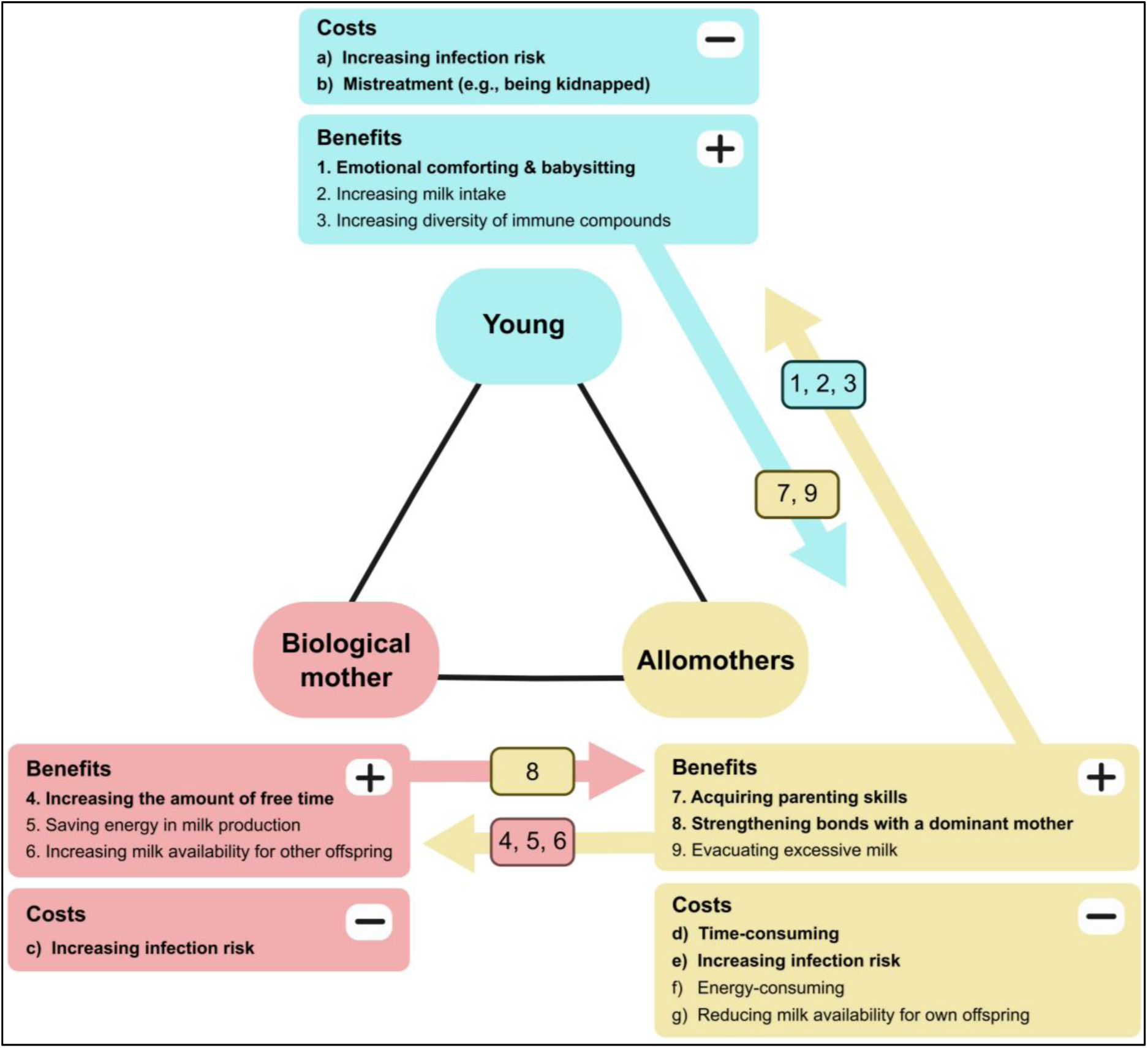
Putative costs (–) and benefits (+) to the three parties (young, allomothers, mother) involved in allonursing. Costs and benefits that are independent of milk transfer from the allomother to the young are in bold. Arrows indicate the transaction of benefits from one party to the other. The numbers inside the arrows indicate the type of benefit given to that party.

**Figure 2.**
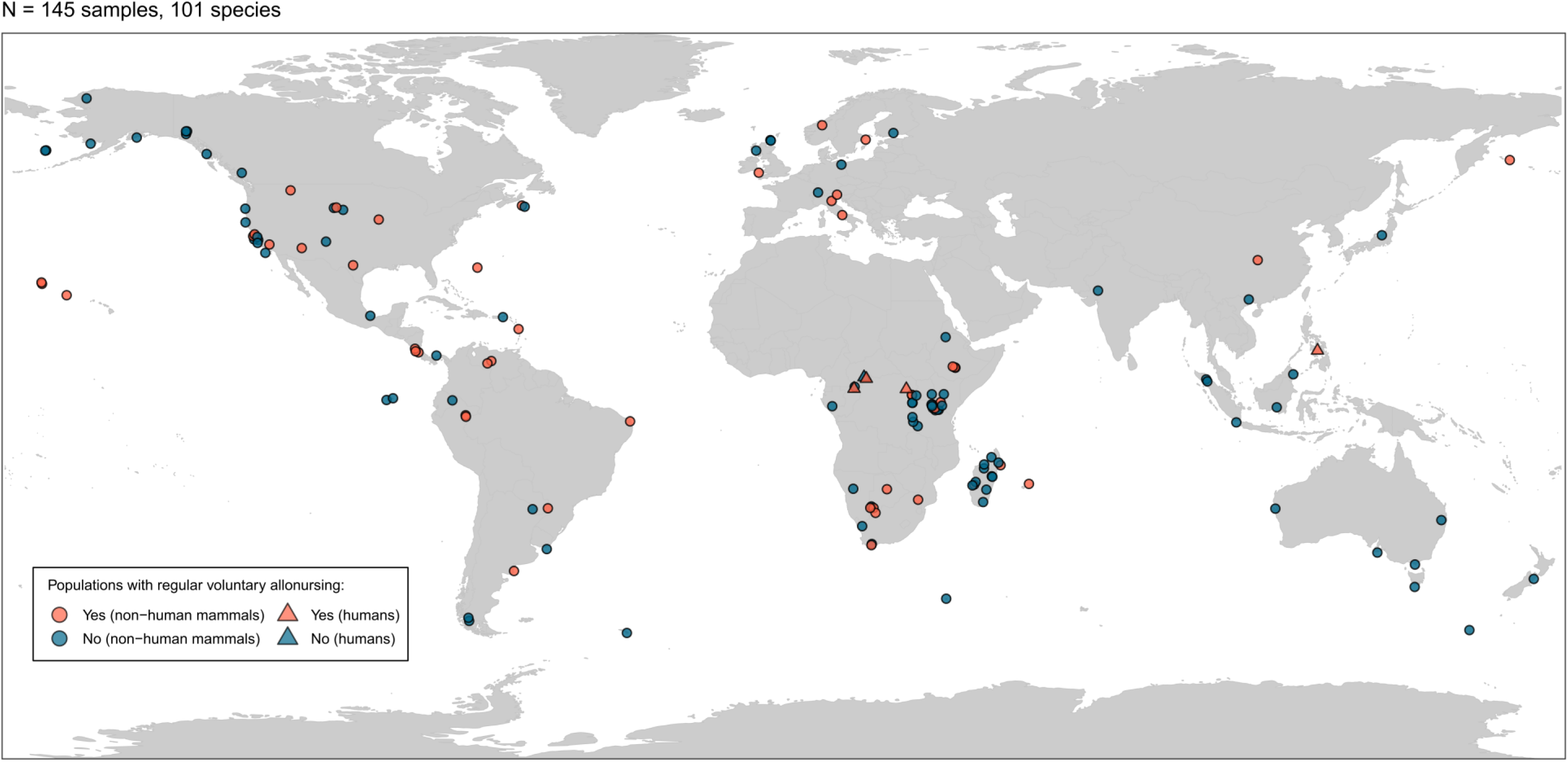
The geographical distribution of samples included in the Co-BreeD *AlloNursing* dataset. Each shape represents a sample from a specific sampling location and sampling period. Circles were slightly jittered (0.5*0.5) to present overlapping sampling locations.

Here, we advance comparative research on allonursing by complementing the Cooperative Breeding Database (Ben Mocha et al., 2025) (hereafter ‘Co-BreeD’; Box 1) with an allonursing dataset. The first part of this paper presents a methodological account of the Co-BreeD *AlloNursing* dataset. Its second part analyses the *AlloNursing* dataset to: (i) map the taxonomic distribution of allonursing, and thereby identify additional mammal species that should be recognized as cooperative breeders; (ii) differentiate cases of non-voluntary from voluntary allonursing to examine to what degree “milk stealing” explains allonursing across species; (iii) quantify the extent of allonursing received by young to evaluate its importance for young and mothers; and (iv) assess the claim that allonursing evolved particularly in species with larger litter sizes (Packer, Lewis & Pusey, 1992; Cerrito & Spear, 2022). The third part of this paper summarises the evidence for the proximate functions of allonursing and articulates further testable predictions for each of these functions.

| **Glossary** |  |
| --- | --- |
| **Cooperative breeding** | **Definition:** A reproductive system where >5% of litters in at least one population receive (i) species-typical parental care from the biological parent/s, and (ii) proactive alloparental care from conspecifics. Alloparental care should fulfil >5% of at least one type of the offspring's needs (Ben Mocha *et al.*, 2023b).<br>**Remarks:**<br>• This definition follows the majority of scholars in viewing cooperative breeding as synonymous with the systematic occurrence of alloparental care (Stacey & Koenig, 1990; Cockburn, 2006).<br>• This definition enables distinguishing the mating system from the care system of a species by considering cooperative breeding regardless of the reproductive status of alloparents (Kappeler, 2019). Thus, it does not differentiate between cooperative and communal breeding species (Ben Mocha *et al.*, 2023a). |
| **Allonursing** | **Definition:** The suckling of a female's nipples by the dependent young of another female.<br>**Remarks:**<br>• This definition does not require retrieving milk from the female or the nursing female lactating since milk retrieval is difficult to infer in field studies (Lappan, 2009; Hewlett & Winn, 2014), and some of the costs and benefits of allonursing can occur without milk retrieval (Figure 1).<br>• This definition excludes the rare cases of suckling from lactating females by nutritionally independent young and adults (O'Brien, 1988). |

### Box 1. The Cooperative Breeding Database (Co-BreeD)

Co-BreeD (Ben Mocha et al.2025) is a database consisting of multiple datasets, each comprising a key biological parameter that is relevant for cooperative breeding research in birds and mammals (including humans). Biological estimates are either: (a) extracted from the primary literature and reviewed by two experts before being sent for verification to an author of the original publication (whenever possible); or (b) obtained as raw unpublished data from researchers and reviewed by two experts.

Co-BreeD has a sample-based structure: each row represents a sample where the biological estimates that were extracted from the sample (e.g., prevalence of breeding events with alloparents and prevalence of allonursing) are linked to the specific sampling location and period. It therefore often includes multiple samples per population and species.

As of today, Co-BreeD includes two complementary datasets that are connected by the *Sample MetaData* section. The *Sample MetaData* columns, each ending with the suffix “_SMD”, provide sample-specific information that is shared by all datasets (e.g., species names, sampling location and period). The first dataset includes a cluster of columns about the *Prevalence of breeding events with potential Alloparents* (“_PA” suffix). The *AlloNursing* dataset consists of the columns with the suffix “_AN”.

Co-BreeD is provided as an R file that is periodically corrected and expanded. The project files are deposited in the open-access Zenodo repository (Ben Mocha (2025) https://zenodo.org/records/14697198). Users are advised to consult Ben Mocha et al. (2025) for a detailed methodological account and recommendations for adequate use.

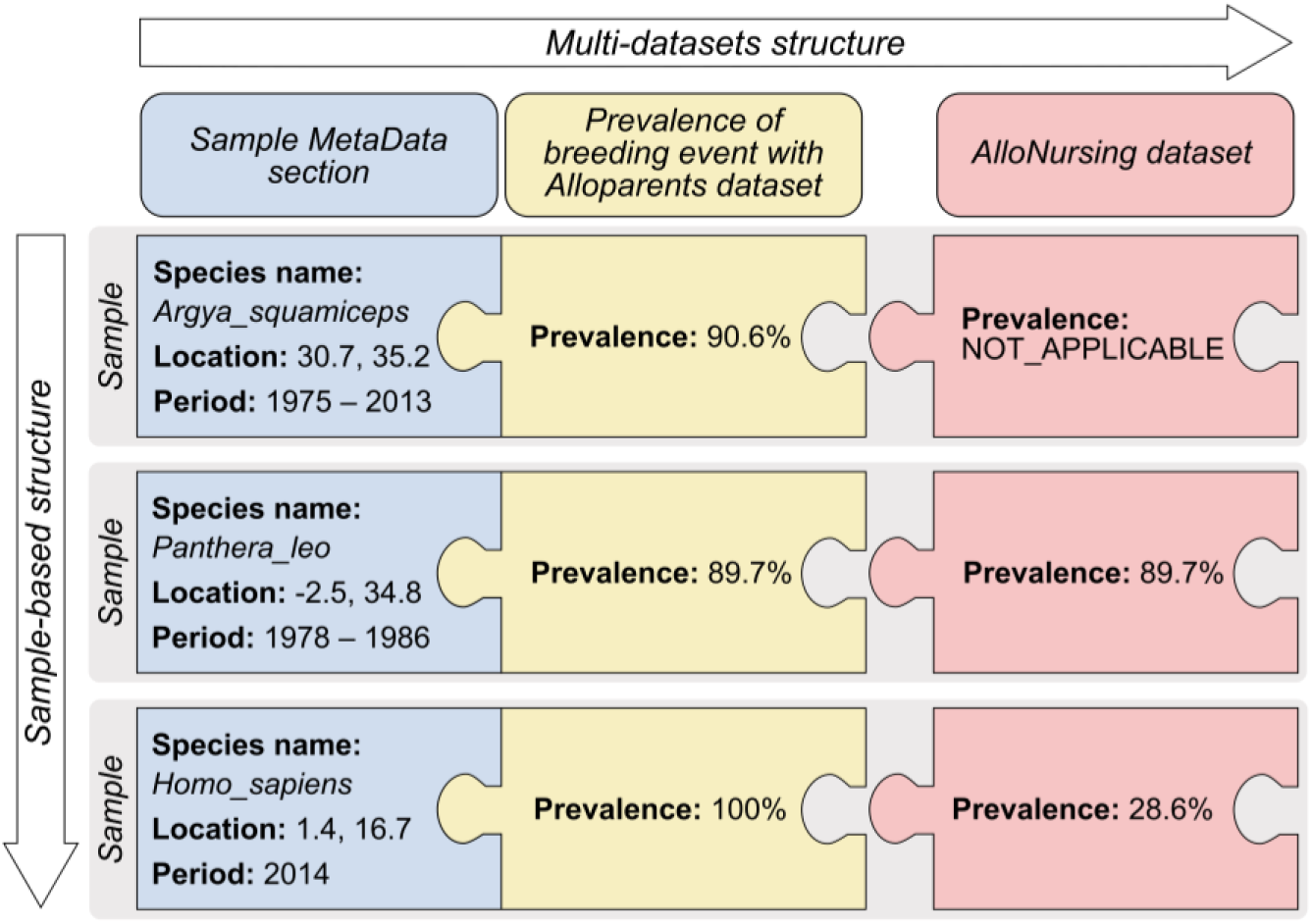

## II. Co-BreeD *AlloNursing* dataset: Allonursing

### II.1 Data collection

Relevant natural history data were gathered in three complementary ways. First, we reviewed all 215 mammal samples included in Co-BreeD V.1 (Ben Mocha et al., 2025). Second, we systematically reviewed all references supporting the classification of the allonursing status of species included in previous allonursing datasets (Packer et al., 1992; MacLeod & Lukas, 2014; Vodenková, 2018) and parental care datasets (Riedman, 1982; Mitani & Watts, 1997; Lukas & Clutton-Brock, 2013; Rosenbaum & Gettler, 2018; Federico et al., 2020). Third, to facilitate representation of taxonomic groups that are under-represented in the current datasets, we (i) invited experts of such species to submit their data, and (ii) searched Web of Science, Google Scholar, and ResearchGate for different taxon names together with “nursing” or “lactation”. To reduce bias towards studies published in English (Amano et al., 2023), Co-BreeD curators repeated these targeted searches and reviewed relevant literature in their native languages (French, Italian, German, Estonian, and Hebrew). This rarely resulted in identifying additional data (N = 2 references).

Our inclusion criteria for studies differed from previous allonursing datasets in three ways. First, we only considered peer-reviewed empirical data. Unpublished data were only included if the methods used to collect them were published elsewhere and the data themselves were reviewed by Co-BreeD curators and published in the Co-BreeD. Second, we only considered data in the primary literature describing wild populations, i.e., species living in a natural habitat with no or minimal human management (e.g., national parks without food supplementation). Data from experimental studies and from captive or domestic animals (including feral populations) were excluded, as these conditions can induce allonursing that is otherwise absent in wild populations (Packer et al., 1992; Kelly, Freeman & Rose, 2025). Third, we only included studies that examined nursing behaviour to ensure that allonursing could have been detected if it occurred. Exceptions to this role were species where allonursing could not occur (e.g., when there was only a single female in all social groups or when infants only interacted with their mothers until weaning).

### II.2. Curated allonursing parameters

#### Categorial occurrence of allonursing

For each sample, the occurrence of allonursing was classified with one of the six categories presented in Table 1.

**Table 1.** Classification categories for the occurrence of allonursing in a sample.

|  |  |
| --- | --- |
| Category | Definition |
| Yes | **Voluntary allonursing was regularly observed in this sample.**<br>• **Voluntary allonursing:** the female was aware of being suckled (e.g., not asleep, looked at the infant) and did not resist it (e.g., by holding the young rather than pushing it away).<br>• **Regular:** allonursing was observed in $\geq 5\%$ of litters (or when we indicate a reason to suspect that the quantitative data is an underestimation). A minimum threshold is needed to avoid rare and potentially exceptional allonursing cases (e.g., North American red squirrel *Tamiasciurus hudsonicus* (Gorrell *et al.*, 2010), eastern grey kangaroo *Macropus giganteus* (King *et al.*, 2015)), disproportionally affecting the classification of the entire sample as exhibiting voluntary allonursing (Griesser & Suzuki, 2016b; Ben Mocha *et al.*, 2025).<br>• Due to paucity of quantitative data, we often relied on the overall verbal descriptions in the text to assess whether $\geq 5\%$ of litters received voluntary allonursing. For instance, “allosuckling is a common behaviour in these species” (Scornavacca *et al.*, 2018, p. 8), “adult females made no discernible attempt to determine the identities of juveniles that nursed from them” (Byers & Bekoff, 1981, p. 779).<br>• Direct observation of suckling was not a necessary condition to infer nursing or allonursing, and indirect evidence of nursing behaviour was sometimes used. For example, milk transfer was inferred via radioactive markers in black-tailed prairie dogs *Cynomys ludovicianus* (Hoogland, Tamarin & Levy, 1989) or specific diving position in sperm whales (Konrad, 2017). |
| Suspected_yes * | **Anecdotal or circumstantial evidence for voluntary allonursing.**<br>• For example, multiple females were observed to nurse multiple offspring in a shared den, but the maternity of each pup was unknown (e.g., (White, 1992)). |
| **No** | **Voluntary and non-voluntary allonursing were not or only rarely observed (<5% of litters).**<br>• “No” was only used when nursing by mothers was examined, so that allonursing could have been detected if it occurred (e.g., Nowell & Fletcher, 2007), or when allonursing could not occurred (e.g., when there was only a single female in all social groups or when infants only interacted with their mothers until weaning (de la Torre *et al.*, 2021)).<br>• If nursing was examined in a study and allonursing was not mentioned, we assumed allonursing did not occur. |
| **Suspected_no \*** | **Nursing was discussed in the publication in a way that suggests regular voluntary allonursing does not occur.**<br>• For instance, the authors considered allonursing was absent in the species with such confidence that they used the lactation status of a female to infer her maternity relationship with the suckling young (e.g., Braude, 1991). |
| **Non_voluntary** | **Young were frequently observed to attempt suckling from females other than their mothers, but were often rejected, resulting in negligible rates of successful allonursing (i.e., <10%).** |
| **MISSING_DATA** | **The study did not examine nursing; thus, allonursing could not be detected.**<br>• These samples are included in Co-BreeD because they provide information for other Co-BreeD datasets.<br>• These samples are not considered for the sample size of the *AlloNursing dataset*. |
| * These samples are included in the *AlloNursing* dataset to inform future research, but due to their uncertainty, they are not considered for analyses in the Results and Discussion section of this paper (N = 17). |  |

#### Prevalence of voluntary allonursing

This parameter estimates the percentage of litters that received voluntary allonursing (i.e., at least one offspring in a litter) out of the total number of litters observed in the sample. Since this parameter was also meant to distinguish between samples with frequent versus anecdotal allonursing, all observed litters were considered regardless of whether allonursing could occur in a litter (i.e., whether there were multiple females in the group or not).

#### Extent of voluntary allonursing

This parameter quantifies the importance of voluntary allonursing for the allonursed offspring. It estimates the percentage of voluntary allonursing bouts (or allonursing time) out of the total number of nursing and voluntary allonursing bouts (or time) 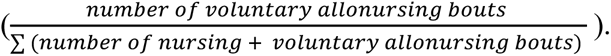 Only offspring that received at least one voluntary allonursing bout were considered. We also categorised whether allonursing bouts were of a shorter/equal/longer duration than nursing bouts (allonursing_vs_nursing_duration_AN column).

### II.3. Hypothesis testing

#### II.3.a. Offspring recognition and rejection

We tested the hypothesis that females allonurse non-voluntarily because they fail to distinguish their offspring from others’ offspring and/or because they cannot reject suckling attempts of young (e.g., if only being suckled when asleep (Packer et al., 1992; Cameron et al., 1999)). To this end, we classified whether (i) the species exhibits evidence that mothers can recognise their offspring (e.g., by exhibiting selective nursing towards specific young (Trillmich, 1981; Dowell, 2005)) (yes/no) and (ii) whether nursing females demonstrated a physical ability to reject suckling attempts (yes/no).

#### II.3.b. Allonursing and litter size

Previous studies reported a positive correlation between allonursing and litter size (Packer et al., 1992; MacLeod & Lukas, 2014; Cerrito & Spear, 2022). The data used in these studies, however, are biased since most species were classified as exhibiting/lacking allonursing based on non-systematic data (e.g., a survey in which experts provided quantitative evaluation based on personal observations (Packer et al., 1992; Cerrito & Spear, 2022)), or a lack of allonursing was inferred based on absence of positive evidence even when nursing behaviour was not studied (MacLeod & Lukas, 2014; Cerrito & Spear, 2022). The latter approach would result in type-two error because if *nursing* behaviour is not studied, *allonursing* is not likely to be detected even if occurring (MacLeod & Lukas, 2014). While these previous methods are informative for obtaining an initial overview and directing future systematic research, their inaccuracy (Schradin, 2017; Ben Mocha et al., 2023b) has led behavioural ecologists (Griesser & Suzuki, 2016; Schradin, 2017) and statisticians (Meng, 2018; Bradley et al., 2021) to recommend against using such data for large-scale comparative analyses. Thus, we revisited the association between allonursing and litter size with four major improvements: (i) using the *AlloNursing* dataset that includes only samples where the presence/absence of allonursing was reliably inferred in wild populations; (ii) accounting for within-species variation by including multiple populations per species; (iii) using allonursing and litter size data from the same population; and (iv) accounting for the degree to which studies underestimate litter size at birth (e.g., fossorial species when litter size could only be recorded after the pups emerged from underground burrow).

#### II.3.b.1. Statistical analyses

Statistical analyses were conducted using R version 4.5.1 (R Core Team, 2022). We fitted two models using a Markov Chain Monte Carlo mixed modelling approach and the MCMCglmm package (Hadfield, 2010). Each model was subjected to 3,000,000 iterations, discarding the initial 100,000 as burn-in and drawing samples every 500 iterations. We detected no evidence of autocorrelation.

To control for phylogeny, we pruned the complete (5,911 species) tip-dated mammal phylogeny from Upham et al. (2019) (https://data.vertlife.org/) to include only the species with relevant data per model. We randomly selected one phylogenetic tree, confirmed its ultrametricity, and used the phylogenetic variance-covariance matrix. For the phylogenetic random effect, we used a weakly informative parameter-expanded inverse-Wishart prior (V = 1, ν = 2; αμ = 0, αV = 25), which permits broad uncertainty in variance estimates while aiding MCMC mixing.

##### Litter size explains allonursing

The first model tested whether monotocous (litter size = 1) and polytocous (litter size >1) species differ in their probability of engaging in voluntary allonursing (allonursing present/absent). To account for within-species variation, the response variable was the proportion of populations exhibiting allonursing out of the number of the species’ populations examined (family = multinomial2). For instance, a species where 3 populations exhibited allonursing and 1 population did not was scored ¾. The model had two predictors: (i) whether the species was monotocous or polytocous, and (ii) whether multiple adult females were associated with each other during the breeding season (yes/no, to account for the possibility of having a potential allomother nearby). There was no within-species variation with regard to this litter size category and female grouping. The effective sample sizes were 2,350 for predictor (i) and 3,030 for predictor (ii).

To furthermore test this explanation for allonursing, we also classified whether allonursing was primarily associated with mothers who lost their offspring (bereaved_mother_AN column: yes/no).

##### Allonursing explains litter size

To test the hypothesis that allonursing enables mothers to produce larger litters, we constructed a model with a Gaussian family. The mean litter size in the population was scaled and set as the response variable (using the scale R function, which subtracts each individual data point from the overall mean and divides it by the standard deviation). The model had two predictors: (i) whether the population exhibited voluntary allonursing (yes/no), and (ii) whether the reported litter size is expected to underestimate litter size at birth (i.e., litter size was measured at birth / at a later phase). To control for multiple populations of the same species, the model had the species ID (independent of phylogeny) as an additional random effect to phylogeny. This random term had a similar prior to the one used for phylogeny (Brouwer & Griffith, 2019). The effective sample sizes were 5,800 for predictor (i) and 6,199 for predictor (ii).

To furthermore test this explanation for litter size, we also classified whether allonursing was primarily directed towards the offspring of a single mother in the social group, who can then save maternal energy without allocating milk to others’ offspring, or whether allonursing was mostly multidirectional (uni_multi_directional_ANcolumn: unidirectional/multidirectional).

For both models, we specified weakly to moderately informative priors. On the residual variance scale, the latent residual variance was fixed at 1 (for the litter size explains allonursing model) or nu = 0.002 (for the allonursing explains litter size model). For the fixed effects, we specified a multivariate normal prior with a mean of zero and a variance–covariance matrix 25*I*, providing moderately informative shrinkage while not strongly constraining parameter estimates. All fixed-effect predictors were mean-centred, so each individual value represents deviations from the overall mean (i.e., each variable was recoded to numeric form, and each individual data point was subtracted from the mean). Data about litter size and the sociality of females during breeding are provided in Table S1.

#### II.3.c. The proximate functions of allonursing

We quantified the empirical support available for the different functions of allonursing (Table 2) by counting the number of samples presenting positive or negative evidence for each function. Given the paucity of quantitative data and explicit hypothesis testing in allonursing research, anecdotal evidence was also considered. For example, cases involving infants of different primate species who survived several days of separation from their mother due to allonursing were considered as evidence that allonursing provides nutrients to young (Perry, 1996; Ren et al., 2012).

**Table 2.**
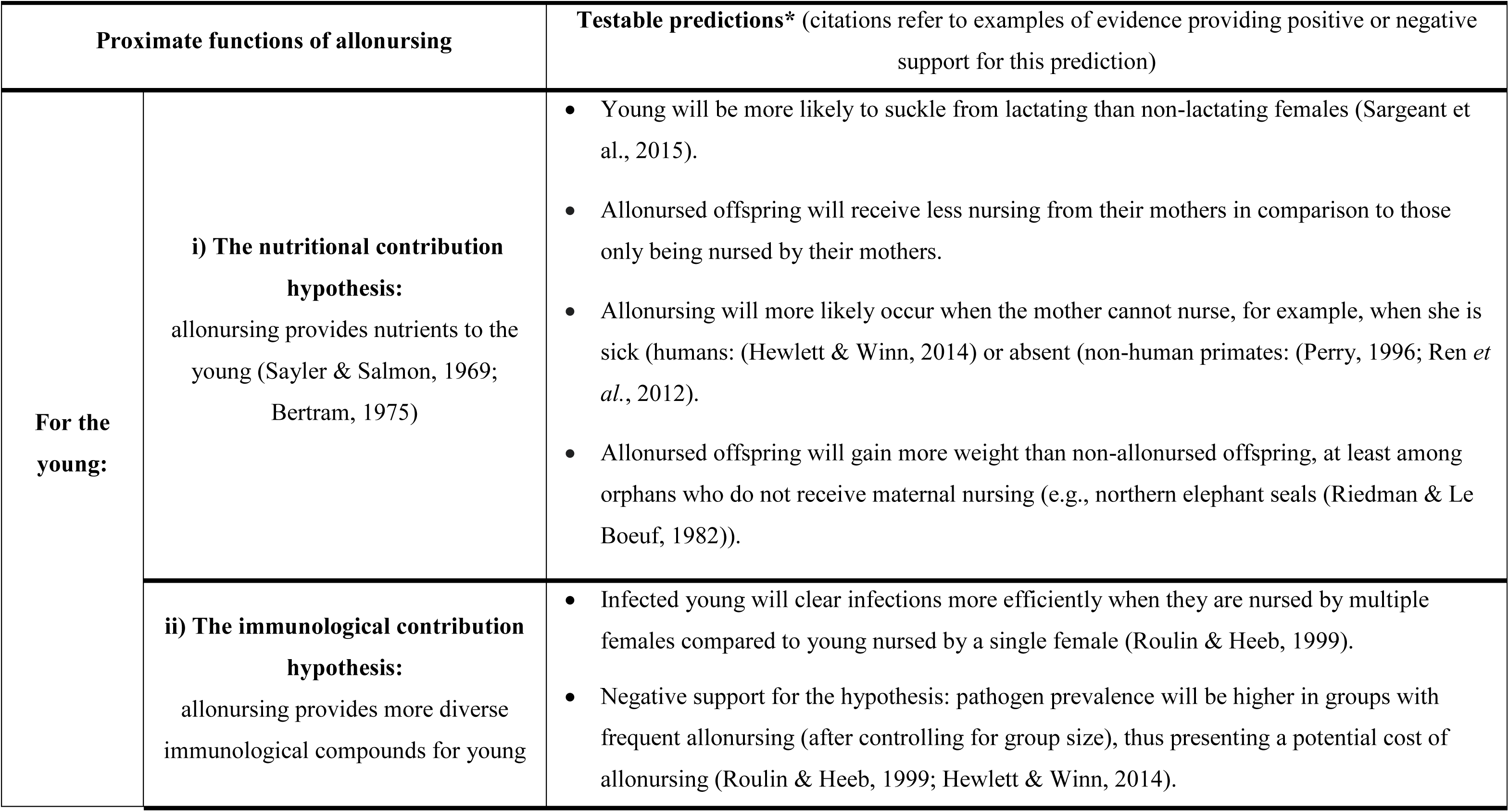
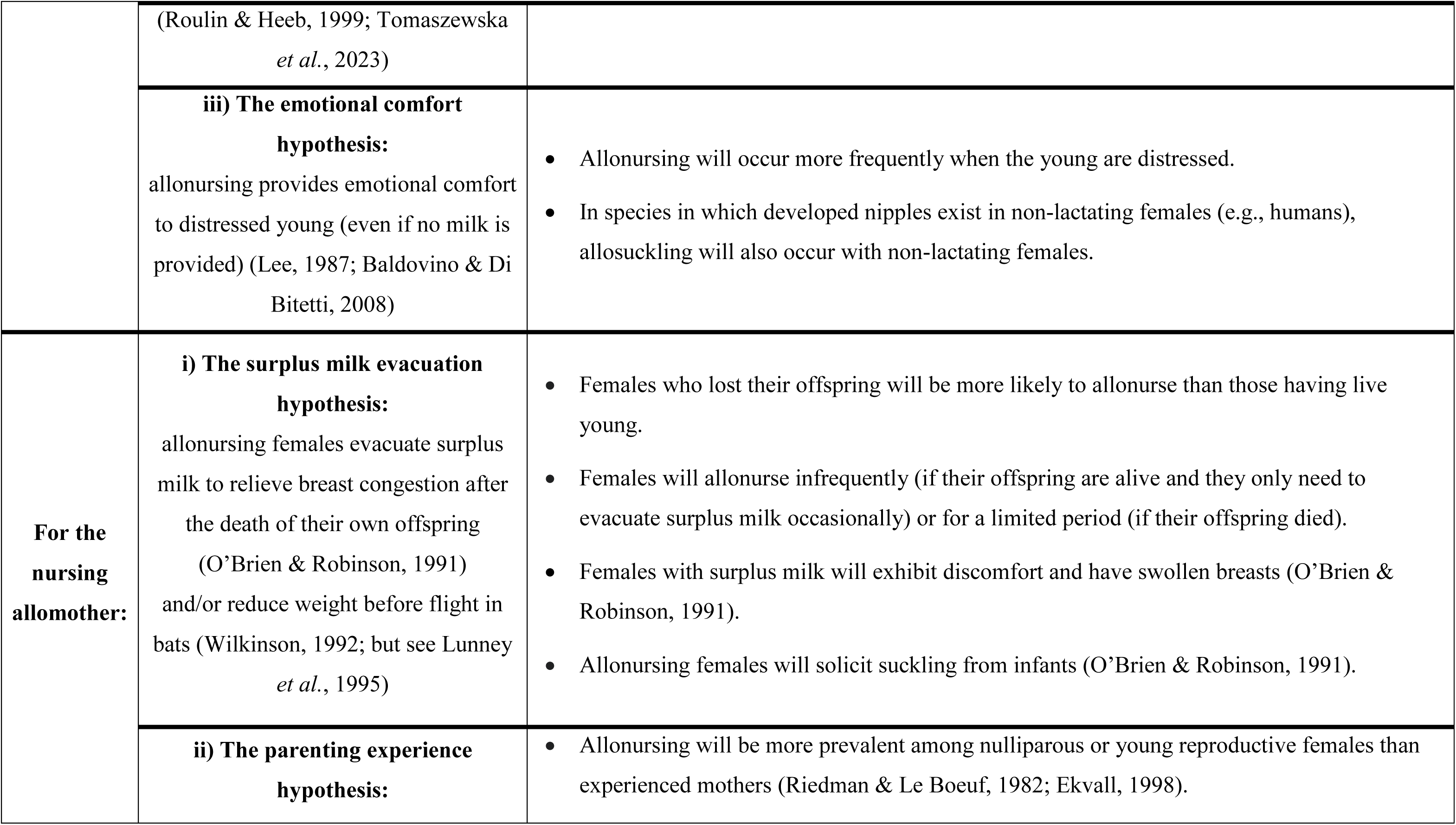
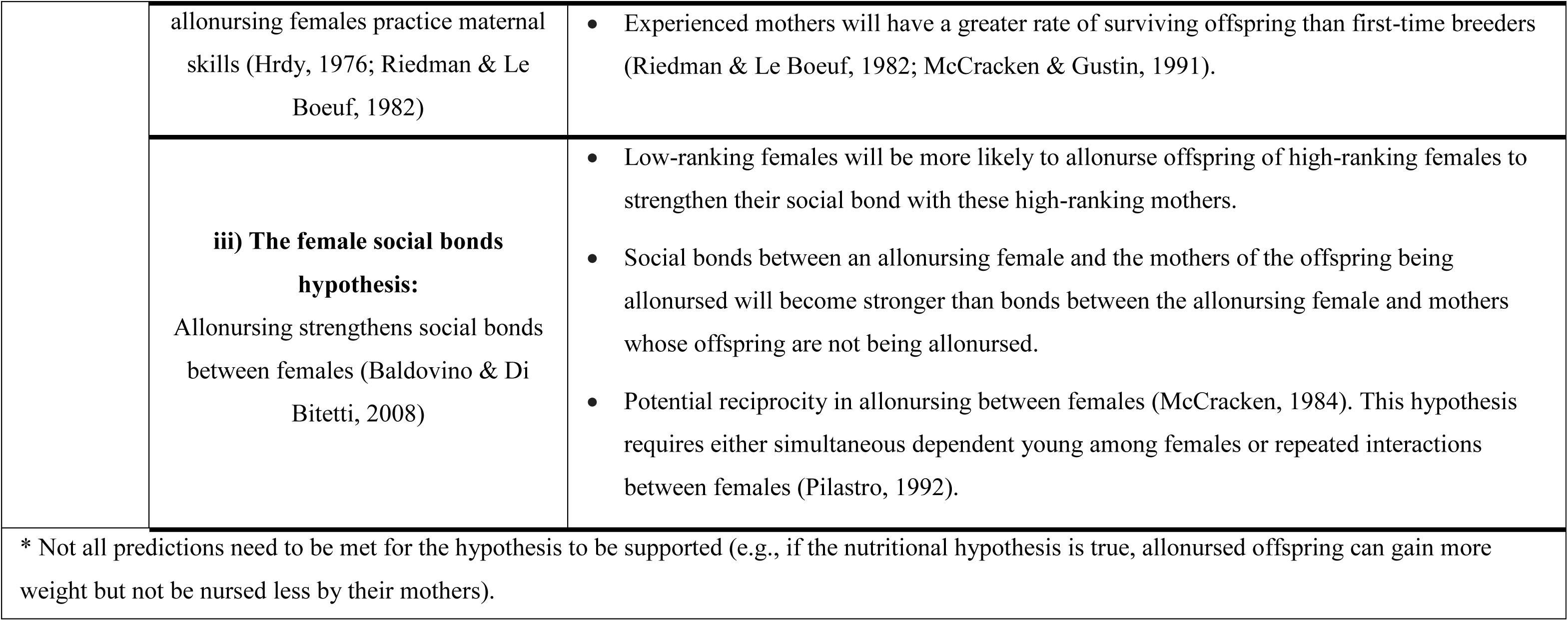
Summary of six non-mutually exclusive hypotheses about the proximate functions of allonursing and their testable predictions.

## III. Results and Discussion

### III.1. Sample characteristics

The *AlloNursing dataset* provides complementary data for the mammal samples included in Co-BreeD V1, and adds 48 new mammal samples to Co-BreeD V2. 145 of these 262 total mammal samples include the minimum categorial data about the occurrence of allonursing and were thus used in the following analyses (i.e., non-voluntary or voluntary allonursing does not occur/occurs, see Table 3).

**Table 3.** Sample size of the Co-BreeD *AlloNursing* dataset (as of December 2025).

|  |  |  |  |  |
| --- | --- | --- | --- | --- |
|  | Number of species | Number of populations | Number of Samples | Samples verified by the author of the original study |
| Non-human mammals | 100 | 129 | 140 | 24% |
| Humans | 1 | 4 | 5 | 60% |
| **Total** | 101 | 133 | 145 | 25%* |
| * An additional 25% of samples could not be verified, and 50% of samples are in the process of verification. |  |  |  |  |

The *AlloNursing dataset* is based on >9,871 litters (not including litters in studies without quantitative data on sample size, see Table 1). Geographically, samples are distributed across the six continents and the Antarctic convergence (Figure 1). Future research should increase representation of allonursing samples from Asia, North Africa, and polar areas, to better account for environmental parameters and taxonomic diversity that characterise these areas and could affect allonursing (Packer et al., 1992; Brouwer & Griffith, 2019).

### III.2. Taxonomic distribution of allonursing

In 36% of the 101 species in the *AlloNursing* dataset, at least one population exhibited voluntary allonursing. These allonursing species are distributed across 5 orders (Artiodactyla, Carnivora, Chiroptera, Primates, Rodentia) out of the 10 orders represented in the *AlloNursing* dataset (orders for which there is currently no evidence of regular voluntary allonursing: Diprotodontia, Hyracoidea, Monotremata, Perissodactyla, Proboscidea). But note the small number of species sampled in these orders that lack allonursing (Figure 3).

**Figure 3.**
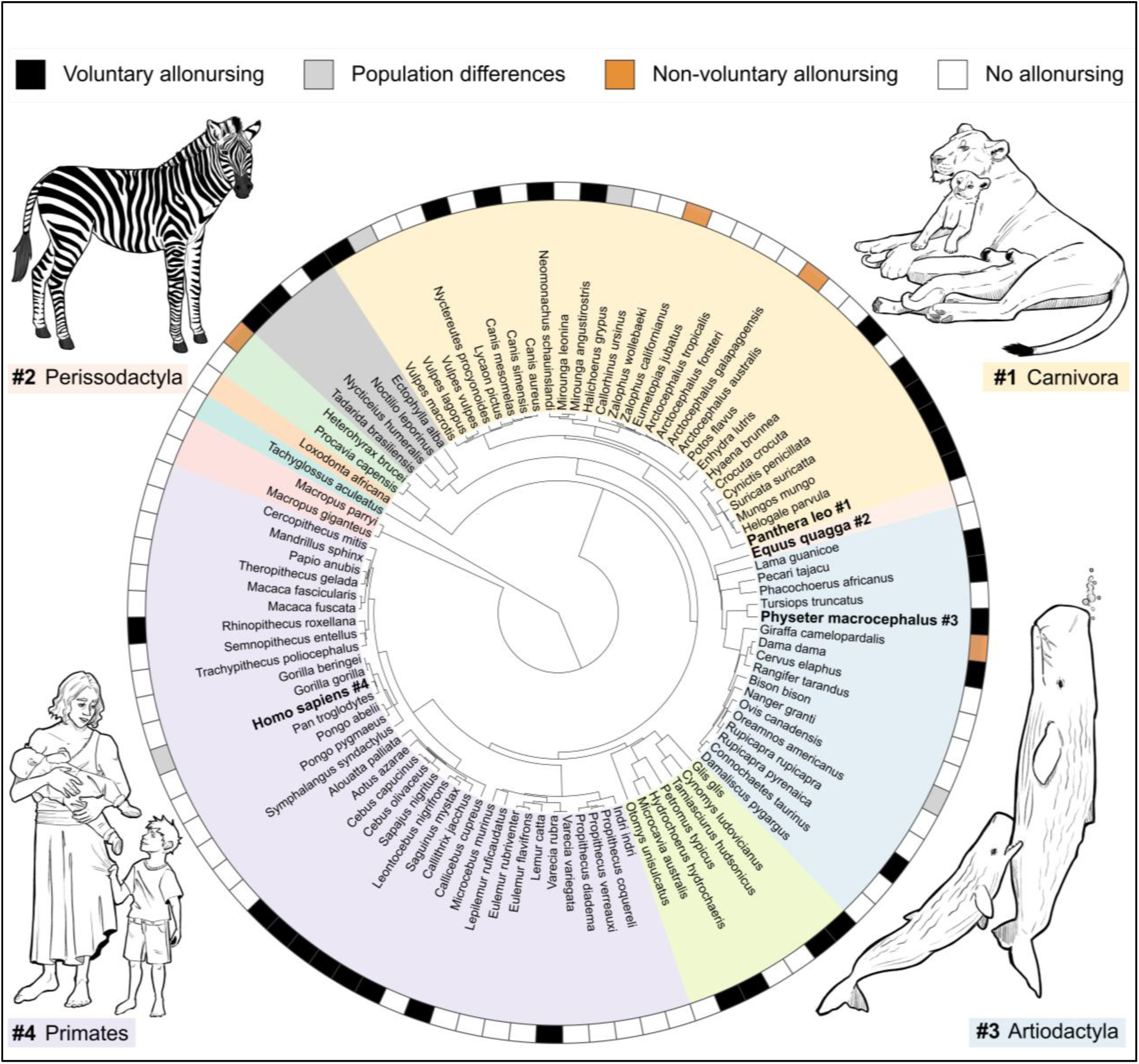
Phylogenetic distribution of voluntary, non-voluntary, and no allonursing across the 10 mammal orders represented in the *AlloNursing* dataset. Taxonomic orders are marked by different colours. Phylogenetic trees were made in phyloT and visualised using iTOL (Letunic & Bork, 2024). Drawn species are marked in bold. Animal illustrations: Natalie Kestel.

Allonursing exhibited moderate intraspecific variability. Four (18%) of the 22 species with data from multiple populations exhibited allonursing in some, but not all, populations studied: Arctic fox *Vulpes lagopus* (allonursing in 2 out of 3 populations), gray seal *Halichoerus grypus* (allonursing in 2 out of 4 populations), bighorn sheep *Ovis canadensis* (allonursing in 1 out of 2 populations), and human *Homo sapiens* (allonursing in 4 out of 5 populations). This intraspecific variability emphasises the importance of using population rather than species-level data in comparative analyses (Griesser & Suzuki, 2016; Ben Mocha et al., 2025).

### III.3. Can non-voluntary allonursing explain the regular occurrence of allonursing?

In some cases, females nursed non-filial young non-voluntarily. This behaviour was speculated to occur because (i) mothers fail to distinguish their offspring from others’ offspring (Packer et al., 1992; Cameron et al., 1999) and/or (ii) females are unable to reject nursing attempts (e.g., if being suckled while asleep (Boness et al., 1998; Maniscalco et al., 2007) or while suckling her own offspring (Ekvall, 1998; Saito & Idani, 2018)). Non-voluntary allonursing is typically readily identifiable. First, young approach inconspicuously to suckle from alien females (Lunn, 1992; Pusey & Packer, 1994). Second, females conspicuously reject non-filial young once they become aware of being suckled by them (Figure 4 (Dowell, 2005)).

**Figure 4.**
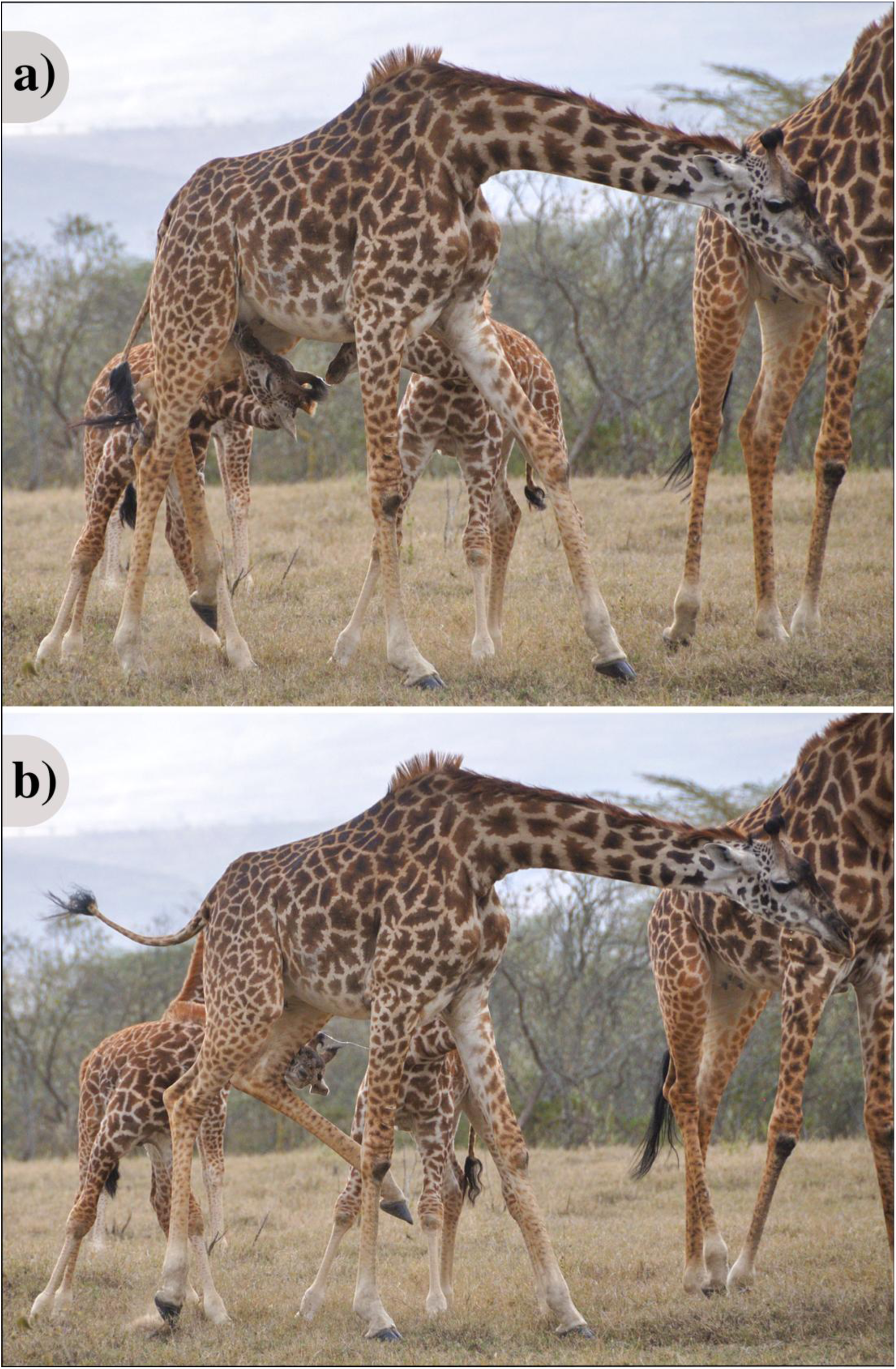
Masai giraffe *Giraffa tippelskirchi* rejects a nursing attempt by a juvenile. Photo: Yitzchak Ben Mocha.

Many of the samples in the *AlloNursing* dataset reported attempts of non-voluntary suckling. However, in only 5 samples, most (≥90%) allonursing attempts were rejected. Consequently, these 5 samples were classified as exhibiting non-voluntary allonursing (e.g., New Zealand fur seal *Arctocephalus forsteri* (Dowell, 2005), Masai giraffe *Giraffa camelopardalis tippelskirchi* (Saito & Idani, 2018)). By contrast, allonursing attempts were accepted in the majority of samples where allonursing was observed (56 out of 61). These 55 samples were therefore classified as exhibiting regular voluntary allonursing.

All 40 species with data on offspring recognition exhibited evidence of mothers recognising their offspring (of which 15 species engaged in voluntary allonursing, 4 in non-voluntary allonursing, and 21 species did not engage in allonursing). Thus, we found no evidence supporting the hypothesis that allonursing usually occurs because mothers fail to recognise their own offspring (Packer et al., 1992; Cameron et al., 1999). Similarly, little evidence was found for the hypothesis that females allonurse due to an inability to reject suckling: in 95% of the 61 species where authors referred to rejection of suckling attempts (of their own or alien offspring), females displayed an ability to reject suckling attempts (of which 19 species engaged in voluntary allonursing, 5 in non-voluntary allonursing and 37 species did not engage in allonursing). The only exceptions were three primate species (gray mouse lemur *Microcebus murinus* (Eberle & Kappeler, 2006), black-horned capuchin *Sapajus nigritus* (Baldovino & Di Bitetti, 2008) and indri *Indri indri* (Weir, 2014)) that exhibited voluntary allonursing, showed high tolerance to infants, and have not been observed rejecting any nursing attempts, though this does not necessarily imply an inability to do so.

Taken together, the *AlloNursing* dataset demonstrates that, although non-voluntary allonursing occurs in many species, it cannot explain the regular occurrence of allonursing in most species, especially where mothers recognise their offspring and females easily reject unwanted nursing attempts (Sargeant et al., 2015).

### III.4. Allonursing and cooperative breeding

According to most scholars (Cockburn, 2006; Kappeler, 2019; review by Ben Mocha et al., 2023b), systematic alloparental care is a necessary and sufficient condition to classify species as cooperative breeders (Glossary). We thus propose that all 33 species exhibiting regular voluntary allonursing and living in cohesive social groups should be classified as cooperative breeders. These include 17 species (e.g., golden snub-nosed monkey *Rhinopithecus roxellana,* kit fox *Vulpes macrotis*, sperm whale) that were not classified as cooperative or communal breeders in previous parental care datasets in mammals (Lukas & Clutton-Brock, 2012; Federico et al., 2020).

### III.5. Allonursing in humans

The *AlloNursing* dataset includes quantitative data for 5 samples from 5 human cultures (Figure 2). Regular allonursing (as defined in Table 1) was reported in four (80%) of these cultures (for qualitative evidence that allonursing occurred in most non-industrial cultures, see (Hewlett & Winn, 2014). This prevalence of allonursing, along with other forms of systematic alloparental care, such as babysitting (Marlowe, 1999; Chaudhary, Salali & Swanepoel, 2023) and food sharing (Hill & Hurtado, 2009) across human cultures, supports the classification of humans as a cooperatively breeding species (Lahdenperä et al., 2004; Sear & Mace, 2008; Burkart, Hrdy & van Schaik, 2009; Hewlett & Winn, 2014; Ben Mocha et al., 2023b). Including humans as a cooperatively breeding species in cross-species comparisons is thus important for integrating research on human evolution into the broader understanding of sociality in the animal kingdom (Hrdy, 2007; Burkart, van Schaik & Griesser, 2017).

### III.6. To what extent does allonursing substitute maternal nursing?

In populations with voluntary allonursing, approximately half of litters received allonursing, although this proportion varied widely across species (median: 44%, range: 3–100%, N = 33 samples from 24 species, Figure 5a). In litters that received allonursing, allonursing bouts often accounted for a relatively modest share of total nursing (median: 25%, range: 7–52%, N = 12 samples from 11 species, Figure 5b). In most species, allonursing bouts were shorter than maternal nursing bouts (64%), and only rarely of similar duration (18%) or longer than maternal nursing (18%, total N = 11 species). We propose that quantitative data about the extent of allonursing (Figure 5b) can be used as a proxy for the extent of time and/or energy saved from biological mothers across species.

**Figure 5.**
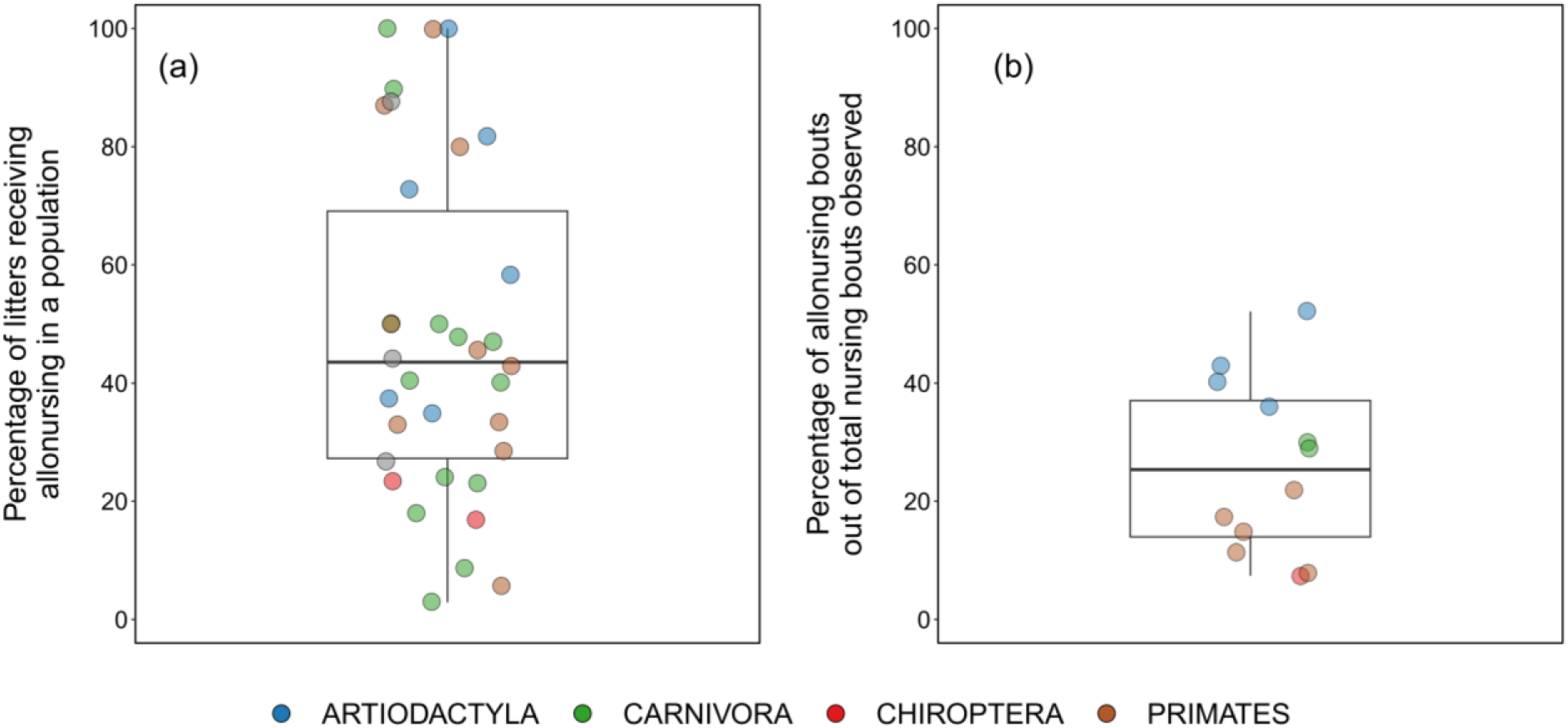
**(a)** The proportion of litters receiving allonursing in populations where regular voluntary allonursing was observed. **(b)** The proportion of allonursing bouts out of total nursing bouts (in litters that received at least one allonursing bout). Horizontal bars represent the median, boxes the 25% and 75% quartiles, and vertical lines indicate the minimum and maximum values.

Occasionally, mothers cannot, permanently or temporarily, nurse their unweaned offspring. For instance, this could occur due to maternal death (common dwarf mongooses *Helogale parvula* (Rood, 1980), North American red squirrels (Gorrell et al., 2010), eastern grey kangaroos (King et al., 2015), pinnipeds (Riedman & Le Boeuf, 1982; Boness et al., 1998; Dowell et al., 2008)), maternal illness (humans (Hewlett & Winn, 2014)), separation of the mother from her social group (primates (Perry, 1996; Ren et al., 2012), bighorn sheep (Hass, 1990)), or maternal abandonment (lions *Panthera leo* (Cairns, 1990)). In some of these cases, allomothers were observed nursing these dependent young until they reunited with their mothers or were weaned (see above references). During prolonged maternal illness or orphaning, allonursing was occasionally documented to have saved the lives of the involved young (see above references). The rarity of such life-saving adoptions, however, resulted in limited systematic investigation into whether they occur more frequently in species that commonly engage in allonursing (e.g., common dwarf mongoose (Rood, 1980)) than in those that do not (e.g., North American red squirrel (Gorrell et al., 2010), eastern grey kangaroo (King et al., 2015)).

Taken together, this evidence indicates that allonursing does not fully replace maternal nursing in any species. Nonetheless, in some species, allonursing constitutes a substantial share of the total nursing received by infants (Figure 5b), and in the relatively rare cases of mother–offspring separation, adopting mothers could substitute for maternal nursing until weaning.

### III.7. Allonursing and litter size

Using the largest population-level dataset on allonursing and litter size curated to date, we revisit two hypotheses below that propose contrasting cause-and-result relationships between allonursing and litter size.

#### Litter size explains allonursing

Packer and colleagues (1992) reported that polytocous species (litter size >1) are more likely to exhibit allonursing than monotocous species (litter size = 1; see also (MacLeod & Lukas, 2014)). We found a similar significant difference: polytocous species were on average ∼42% more likely to engage in allonursing than monotocous species (posterior mean = 3.8, 95% CI = 1.3 to 6.7, pMCMC = 0.002; N = 110 populations, 87 species).

To partly explain this association between allonursing and litter size, Packer and colleagues (1992) suggested the “less-costly allonursing” hypothesis. According to this hypothesis, allonursing is more costly for monotocous mothers because if they lose their single offspring, they have to continue producing milk just for allonursing, while polytocous mothers who lose part of their litter would anyway continue producing milk for their remaining offspring (Packer et al., 1992; Roulin, 2002). Our *AlloNursing* data, however, show that in only 14% of species (total N = 35), voluntary allonursing was primarily associated with females who lost their offspring. Furthermore, while the less costly allonursing hypothesis suggests a reduction of constraints for allonursing, it does not explain why selection would favour females paying any cost to allonurse others’ offspring. We thus propose that the litter size explains little variation in allonursing. Indeed, the widely overlapping credible intervals for the two categories (Figure 6a) suggest that substantial variation in allonursing remains unexplained even after the model accounted for litter size, female sociality during breeding, phylogeny, and population identity.

**Figure 6.**
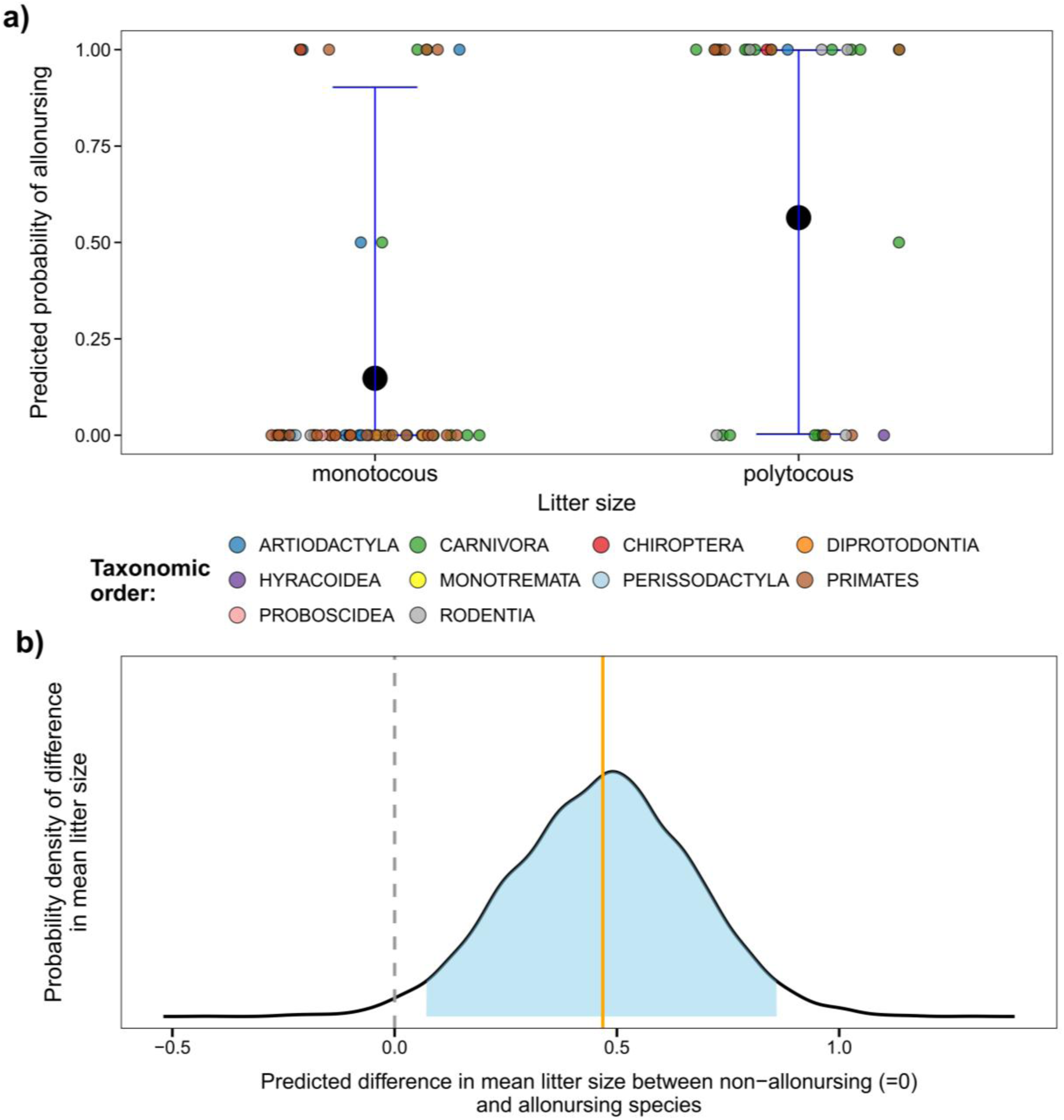
**(a)** Predicted probability of species to engage in voluntary allonursing as a function of litter size: monotocous (litter size = 1) vs polytocous (litter size > 1). Each jittered point shows the observed proportion of populations engaging in allonursing for a given species. The enlarged black points indicate the predicted posterior mean probability from the model, and blue error bars represent the 95% credible intervals of these means (N = 110 population, 87 species). **(b)** The predicted mean difference in litter size between populations that do not exhibit allonursing (0, dashed grey line) and populations that engage in allonursing. The black curve represents the posterior probability distribution of the true mean difference (the orange solid line indicates the mean probability at X = 0.47. The shaded blue area corresponds to the 95% credible intervals of this probability (N = 102 populations, 80 species).

#### Allonursing explains litter size

The “release-of-maternal-energy” hypothesis argues that the nutrients provided to young by allonursing enable their mothers to divert metabolic energy into increasing litter size (Cerrito & Spear, 2022). It thus predicts a positive correlation between litter size and the occurrence of allonursing (see supporting evidence in (Cerrito & Spear, 2022)). Using the *AlloNursing* dataset, we confirmed the positive correlation between mean litter size and allonursing (posterior mean = 0.33, 95% CI = 0.06 to 0.61, pMCMC = 0.021, N = 102 populations, 80 species; Figure 6b). Nonetheless, our model finds considerably smaller effect size (+0.47 young per litter in species exhibiting allonursing, Figure 6b) than in a previous analysis (+1.3 young per litter (Cerrito & Spear, 2022)).

The interpretation that larger litters are the consequence of saved maternal energy (Cerrito & Spear, 2022) inexplicitly assumes that allonursing is unidirectional from allonursing females to specific mothers. Only under this assumption of unidirectional flow of allonursing (i.e., energy), some mothers can redirect saved metabolic energy to increase litter size. The *AlloNursing* data, however, refute this unidirectional assumption: voluntary allonursing was seldom directed towards the offspring of a single female (6 out of 30 species). Rather, in most allonursing species, group members allonursed each other’s offspring simultaneously (albeit not equally), and females therefore did not necessarily save metabolic energy that allowed all of them to produce more young.

#### Allonursing and litter size

In sum, while the *AlloNursing* dataset confirms the positive correlation between allonursing and litter size (Packer et al., 1992; MacLeod & Lukas, 2014; Cerrito & Spear, 2022), the synthesis between its behavioural parameters reveals two important insights. First, the effect of allonursing on litter size is considerably smaller than previously reported, and its biological significance is therefore likely smaller than previously suggested (Cerrito & Spear, 2022). Second, while allonursing could be less costly for polytocous species (Packer et al., 1992; MacLeod & Lukas, 2014) and save some maternal energy (Cerrito & Spear, 2022), our analyses suggest that neither of these hypotheses fully explain the association between allonursing and litter size. We propose instead that allonursing facilitates offspring survival and thereby increases litter size at weaning rather than at birth (see section III.8 on the function of allonursing and also (Packer et al., 1992)).

### III.8. The proximate functions of allonursing

A fundamental question is, “what are the proximate functions of allonursing for the allonursing females and for receiving young?” To facilitate future research, we (i) summarised the six main suggested functions and proposing testable predictions for each hypothesis (Table 2), (ii) quantify the degree of empirical support for each hypothesis (Figure 7), and (iii) map the potential costs and benefits for each party involved in allonursing: the young receiving allonursing, their mother, and the female providing allonursing (Table S2 and Figure 1). Note that the ultimate pressures that select and maintain allonursing − e.g., kin selection or reciprocity − are beyond the scope of this study.

**Figure 7.**
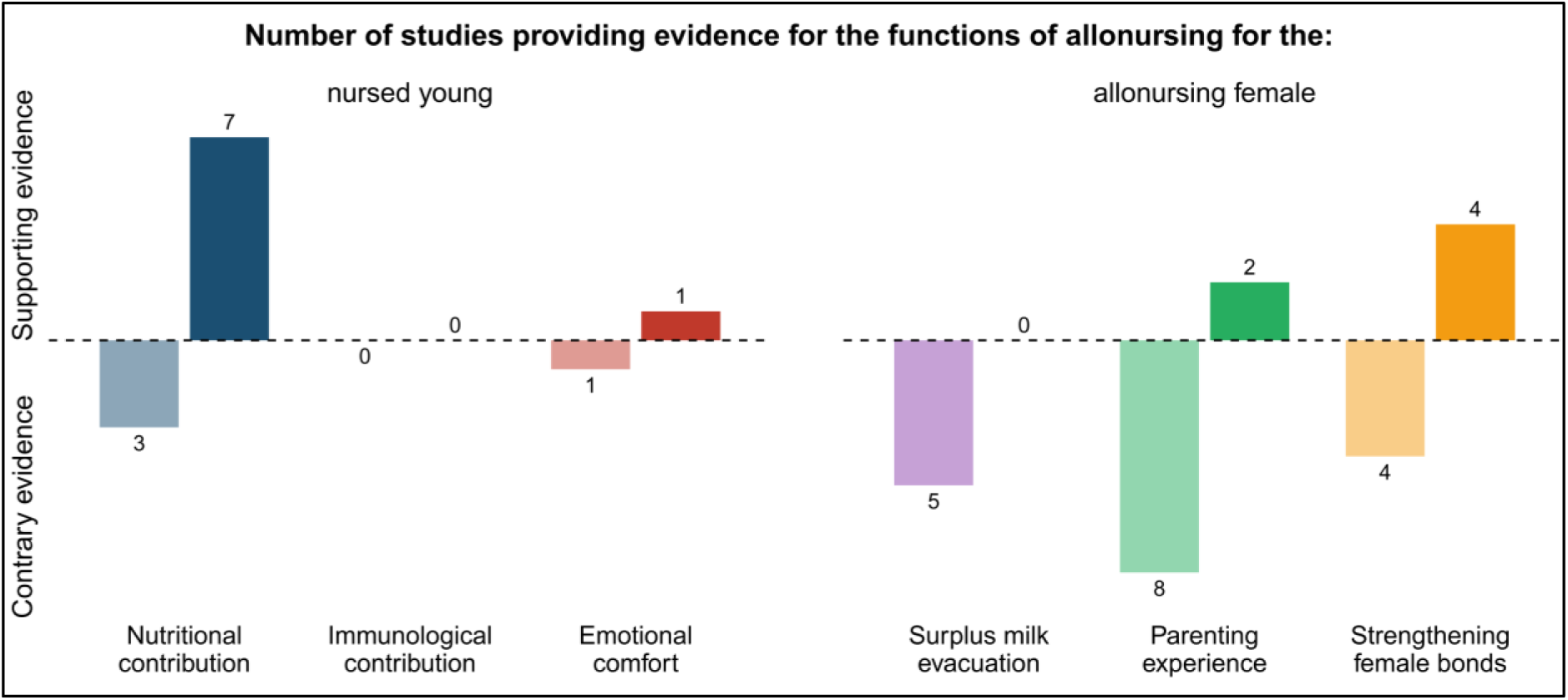
The number of studies that provide at least some supporting or contrary evidence for the main proximate functions of allonursing. (see Table S3 for further information and references).

As shown in Tables 2 and S1, the six major hypotheses are not mutually exclusive, and allonursing can concurrently fulfil several functions within the same population (e.g., providing nutrients, emotional support and diverse immunological compounds for the young, while strengthening the relationships between the mother and allomothers, and providing the latter maternal experience, all at the same time (Sargeant et al., 2015)). This overlap makes it challenging to disentangle different hypotheses. Nonetheless, Figure 7 demonstrates that provisioning is the benefit/function of allonursing that is most commonly received empirical support. We note that this may be a by-product of the methodological difficulties in testing other functions.

We propose distinguishing between allonursing in the absence versus the presence of maternal nursing. When the biological mother is incapable of nursing temporarily (e.g., due to illness or separation from the group) or permanently (e.g., due to her death), the main function of allonursing is provisioning. Breeding in a group with multiple nursing females can, therefore, provide an insurance against the incapability of maternal nursing (Rudnai, 1973; Rood, 1980; Eberle & Kappeler, 2006; Chaudhary et al., 2023). When mothers are capable of nursing their young, allonursing can supply additional milk for them (Riedman & Le Boeuf, 1982; Hoogland et al., 1989; Sarano et al., 2023), while simultaneously freeing up time for mothers through *de facto* babysitting. The proportion of allonursing presented in Figure 5b could be used as an approximation of the proportion of metabolic energy and time saved by mothers. Note that maternal metabolic energy is only saved when allonursing is relatively unidirectional, but since multiple young can be nursed simultaneously, allonursing can save maternal time also when allonursing is reciprocal.

While the nutritional contribution and free time provided by allonursing is sometimes limited (Figure 5b), these gains often join additional benefits gained from other forms of alloparental care to potentially constitute a significant contribution to young and/or mothers. In these cases, measuring the combined effect of alloparental care by comparing the survival of assisted versus unassisted litters is necessary to assess the impact of allonursing and alloparental care in general (Macleod, McGhee & Clutton-Brock, 2015).

## IV. Conclusions

1. The Cooperative-Breeding Database (Co-BreeD) is an open-access and peer-reviewed resource that integrates different datasets, each dedicated to a key biological parameter relevant for cooperative breeding research in birds and mammals (Box 1 (Ben Mocha et al., 2025)). The hereby curated *AlloNursing* dataset complements Co-BreeD with the largest number of samples with systematic data on nursing in wild populations (Table 3).
2. Alloparental care is a sufficient condition to categorise species as cooperative breeders (Cockburn, 2006; Ben Mocha et al., 2023b). The *AlloNursing* dataset expands the list of cooperatively breeding mammals by identifying 17 additional species that exhibit regular voluntary allonursing, but were previously not classified as communal or cooperative breeders in comparative datasets.
3. Cases of non-voluntary allonursing (“milk stealing”) occur in many species. However, because females are often aware of being suckled, actively support allosuckling, and can recognise their offspring, non-voluntary allonursing explains systematic allonursing in only a handful of species.
4. We present the first quantitative data on the prevalence (Figure 5a) and extent (Figure 5b) of allonursing, which can be used to quantify the significance of allonursing for biological mothers across mammals.
5. Phylogenetic mixed models using the *AlloNursing* data reject current explanations for the association between allonursing and larger litter size. Instead, we propose that allonursing may facilitate offspring survival and thereby increase litter size at the weaning stage.
6. Nutrient provisioning is the most empirically supported proximate function of allonursing for the allonursed young (Figure 7). We suggest that allonursing simultaneously frees up time for mothers and can provide young and mothers with an ‘insurance policy’ against maternal death (Perry, 1996; Eberle & Kappeler, 2006; Hewlett & Winn, 2014). Further studies should focus on explaining the benefits to the allonursing females.
7. Co-BreeD is an updatable resource (https://zenodo.org/records/14697198 (Ben Mocha, 2025)). We thus encourage field researchers to contribute data about parental care from wild populations of any bird and mammal species and/or suggest corrections to the database (see ESM data contribution form). Data contributors will be provided with the opportunity to co-author a methodological paper incorporating these updates.

## Authors contribution

**Research design:** Yitzchak Ben Mocha

**Funding acquisition:** Yitzchak Ben Mocha, Michael Griesser

**Project management:** Yitzchak Ben Mocha, Maike Woith

**Data collection:** Yitzchak Ben Mocha, Maike Woith, Sophie Scemama de Gialluly, Lucia Bruscagnin, Laura Pipper, Natalie Kestel

**Data analysis:** Yitzchak Ben Mocha

**Visualisation:** Yitzchak Ben Mocha, Lucia Bruscagnin, Natalie Kestel

**Data validation:** Yitzchak Ben Mocha, Maike Woith, Sophie Scemama de Gialluly, Lucia Bruscagnin, Laura Pipper, Nikhil Chaudhary, Paul Alan Garber, Lee T. Gettler, Christine M. K. Clarke, Andrea Pilastro, Heinz Richner, Stacy Rosenbaum, Rubén Quintana, Eduardo S. A. Santos

**Contribution of year-by-year data:** Heinz Richner, Sonny Agustin Bechayda, Christine M. K Clarke, Lee T. Gettler, Andrea Pilastro, Andrew N. Radford, Stacy Rosenbaum

**Writing of manuscript:** Yitzchak Ben Mocha

**Editing of the manuscript:** Yitzchak Ben Mocha, Maike Woith, Sophie Scemama de Gialluly, Lucia Bruscagnin, Laura Pipper, Natalie Kestel, Nikhil Chaudhary, Paul Alan Garber, Lee T. Gettler, Christine M. K Clarke, Andrea Pilastro, Heinz Richner, Stacy Rosenbaum, Rubén Quintana, Andrew N. Radford, Eduardo S. A. Santos, Michael Griesser

## Acknowledgements

This work was funded by the Deutsche Forschungsgemeinschaft (DFG, German Research Foundation) under Germany’s Excellence Strategy – EXC 2117 – 422037984. Further funding were provided by the ZENiT grant (the Federal Ministry of Education and Research and the Baden-Württemberg Ministry of Science) and the Klaus Tschira Stiftung gGmbH awarded to Yitzchak Ben Mocha. Michael Griesser was supported by a Heisenberg Grant number: R 4650/2-1 by the German Research Foundation DFG.

